# Four large indels detected by cpTILLING in barley chloroplast mutator seedlings

**DOI:** 10.1101/2021.02.19.432034

**Authors:** F. Lencina, A.M. Landau, M.G. Pacheco, K. Kobayashi, A.R. Prina

## Abstract

In a previous work, a polymorphism detection strategy based on mismatch digestion was applied to the chloroplast genome of barley seedlings that carried the chloroplast mutator (*cpm*) genotype through many generations. Sixty-two different one- or two-nucleotide-polymorphisms were detected along with four large indels: an insertion of 15 bp in the intergenic region between *tRNA^His^* and *rps19* genes, a deletion of 620 bp in the *psbA* gene, a deletion of 79 bp in the intergenic region between *rpl33* and *rps18* genes and a deletion of 45 bp in the *rps3* gene. In the present investigation, we analyzed direct repeats located at the borders of those four large indels. Furthermore, *we* investigated the consequences of protein expression of large indels located in coding regions. The deletion of 620 bp in the *psbA* gene was lethal at the second leaf stage when homoplastomic. The deletion of 45 bp in the *rps3* gene, which eliminates 15 amino acids, did not affect the viability of the seedlings in homoplastomy. Interestingly, the deleted segment is also lacking in the wild type version of the *rps3* gene of maize and sorghum. The presence of direct repeats at the borders of the four large indels suggests that they could have originated by illegitimate recombination. This would be in agreement with a previous hypothesis that the *Cpm* gene product would correspond to a mismatch repair (MMR) protein devoted to maintain plastome stability by playing fundamental roles in mismatch repair during replication and avoiding illegitimate recombination.

## Introduction

Some nuclear gene mutants cause an increased frequency of mutations in the highly conserved plastid genome (Kirk and Tilney-Basset, 1986; Börner and Sears, 1986; Prina *et al*., 2012). One of these mutants is the barley chloroplast mutator (*cpm*), which was at first described as inducing a wide range of cytoplasmically inherited chlorophyll mutations (Prina, 1992). A polymorphism detection strategy based on mismatch digestion named cpTILLING (chloroplast Targeting Induced Local Lesions in Genomes), was used to investigate the presence of plastome polymorphisms in 304 *cpm* seedlings. These seedlings corresponded to two groups of families that carried the *cpm* gene during different number of generations of natural auto-pollination (see Landau *et al*., 2016). After the analysis of 31 PCR amplicons comprising 33 genes and some intergenic regions of the plastome, 62 different one- or two-nucleotides polymorphisms were identified, indicating that *cpm* seedlings carried a highly unstable plastome. The vast majority of those polymorphisms were due to substitutions and small indels (insertions/deletions) in microsatellites. However, a peculiar spectrum that included several combinations of five single nucleotide polymorphisms was observed in the *rpl23* gene, which was later explained as arisen not from mutations, but from increased illegitimate recombination between the *rpl23* gene and its pseudogene (Lencina *et al*., 2019). Besides the polymorphisms mentioned above, four large indels were found, which are the subject of analysis of the present work. Short identical sequences *i.e*. perfect direct repeats or short similar sequences *i.e*. imperfect direct repeats were identified at both ends of the four large indels. This result suggests that, as in the case of the polymorphisms observed in the *rpl23* gene and its pseudogene (Lencina *et al*., 2019), they also may be due to increased illegitimate recombination caused by malfunction of the *Cpm* gene product. Furthermore, we investigated the consequences of protein expression of two of the large indels located in coding regions.

## Materials and methods

### Plant material

The four chloroplast mutator (*cpm*) seedlings carrying the four large indels studied here (one different large indel per seedling) were previously identified through a cpTILLING approach (Landau *et al*., 2016). Genomic DNA samples were isolated from *cpm* and wild type (WT) barley seedlings. The latter corresponded to the parental genotype, which was mutagenized and then used to isolate the *cpm* mutant (Prina 1992). Different *cpm* families were obtained after several generations of natural auto-pollination of seedlings coming from crosses WT (female) by *cpmcpm* seedlings (male). Large indels in rps19 (1,274 bp) and psbA (1,390 bp) PCR amplicones were observed in *cpm* seedlings corresponding to the F_12_ generation coming from two different F_3_ plants that were originated in the same F_2_ plant. The large indels in the rpl33 (1,085 bp) and the rps3 (1,334 bp) amplicons were observed in *cpm* seedlings corresponding to the F_6_ generation coming from two different F_2_ plants. The polymorphisms identified were an insertion of 15 bp in the intergenic region between *tRNA^His^* and *rps19* genes in the rps19 amplicon. A deletion of 620 bp in the *psbA* gene in the psbA amplicon, a deletion of 79 bp in the intergenic region between *rpl33* and *rps18* genes in the rpl33 amplicon and a deletion of 45 bp in the *rps3* gene in the rps3 amplicon.

In addition, we analyzed a few seedlings corresponding to the progeny coming from the family in which the *psbA* gene polymorphic plant mentioned above was found.

### DNA isolation

Genomic DNA was isolated from one or two leaves of individual seedlings using the micromethod described by Dellaporta (1994) with modifications. The tissue was ground with Dellaporta isolation buffer in the Fast Prep®-24 Instrument (MP Biomedicals, USA) and extracted with chloroform before DNA precipitation. DNA concentrations were measured using a Nanodrop spectrophotometer (Thermo Scientific, Wilmington, DE, USA) and standardized to a concentration of 80 ng/μl.

### Celery juice extract (CJE) digestions and electrophoresis of amplicons

The four amplicons carrying the large indels mentioned above (see Plant material), were amplified in two ways: using as a template the *cpm* DNA mixed with wild type DNA or using the *cpm* DNA alone. Afterwards, a step of denaturation and slow re-annealing was added to favor the formation of heteroduplexes at the end of the PCR reaction. Finally, the amplicons were digested with CJE (Till *et al*., 2006) and subjected to electrophoresis in 3% non-denaturing polyacrylamide gels as described in Landau *et al*. (2016). The amplicons were also subjected to electrophoresis without performing previous CJE digestion.

### Sequence analysis

DNA sequences of the four loci carrying large indels and the reading frames of the *psbA* and *rps3* genes were compared with the corresponding wild type sequences through alignments with Clustal O (1.2.4). In addition, barley RPS3 protein from wild type and deleted gene sequences were compared with the protein sequences from other grass species.

### Thylakoid membrane isolation and immunoblotting

Thylakoid membranes were isolated according to Guiamet *et al*. (2002). Thylakoid proteins were separated by SDS-PAGE and immunoblotted with several antibodies to determine the presence or absence of different polypeptides. Solubilized thylakoids of each sample corresponding to the same amount of chlorophyll were electrophoretically separated in 13% (w/v) SDS-polyacrylamide gels. Separated proteins were transferred to nitrocellulose membranes and probed with rabbit antibodies raised against A/B, D, E, L photosystem I (PSI) proteins (kindly provided by H.V. Scheller, The Royal Veterinary and Agricultural University, Copenhague, Denmark), LHCB1 photosystem II (PSII) light harvesting complex protein (kindly provided by J.J. Guiamet, INFIVE, Universidad de La Plata, Argentina), D1 and D2 PSII reaction center proteins (kindly provided by Prof. A. Barkan, University of Oregon). Immunodetection was accomplished with the SuperSignal West Dura extended duration substrate (Pierce) according to the manufacturer’s description and using horseradish peroxidase-conjugated goat-anti-rabbit secondary antibody.

## Results

### Four large indels were detected by CJE digestion

The patterns of digestion with celery juice extract (CJE) of the four amplicons carrying large indels (see Materials and Methods) are shown in Fig. 1. In addition to the full-size wild type band and the fragments that result from its digestion, a second band was observed for the rps19, psbA and rpl33 amplicons. These bands were larger than the wild type version of the rps19 amplicon and smaller for those of the psbA and rpl33 amplicons. In contrast, only one amplification band was observed for the rps3 amplicon (Fig. 1). Sequencing of the amplification band from the rps19 amplicon revealed that it contained a 15 bp insertion, while the rps3 amplicon had a deletion of 45 bp. In the psbA and rpl33 amplicons, sequencing of amplification bands purified from agarose gels revealed the existence of deleted versions lacking segments of 620 bp and 79 bp, respectively.

**Figure 1.**
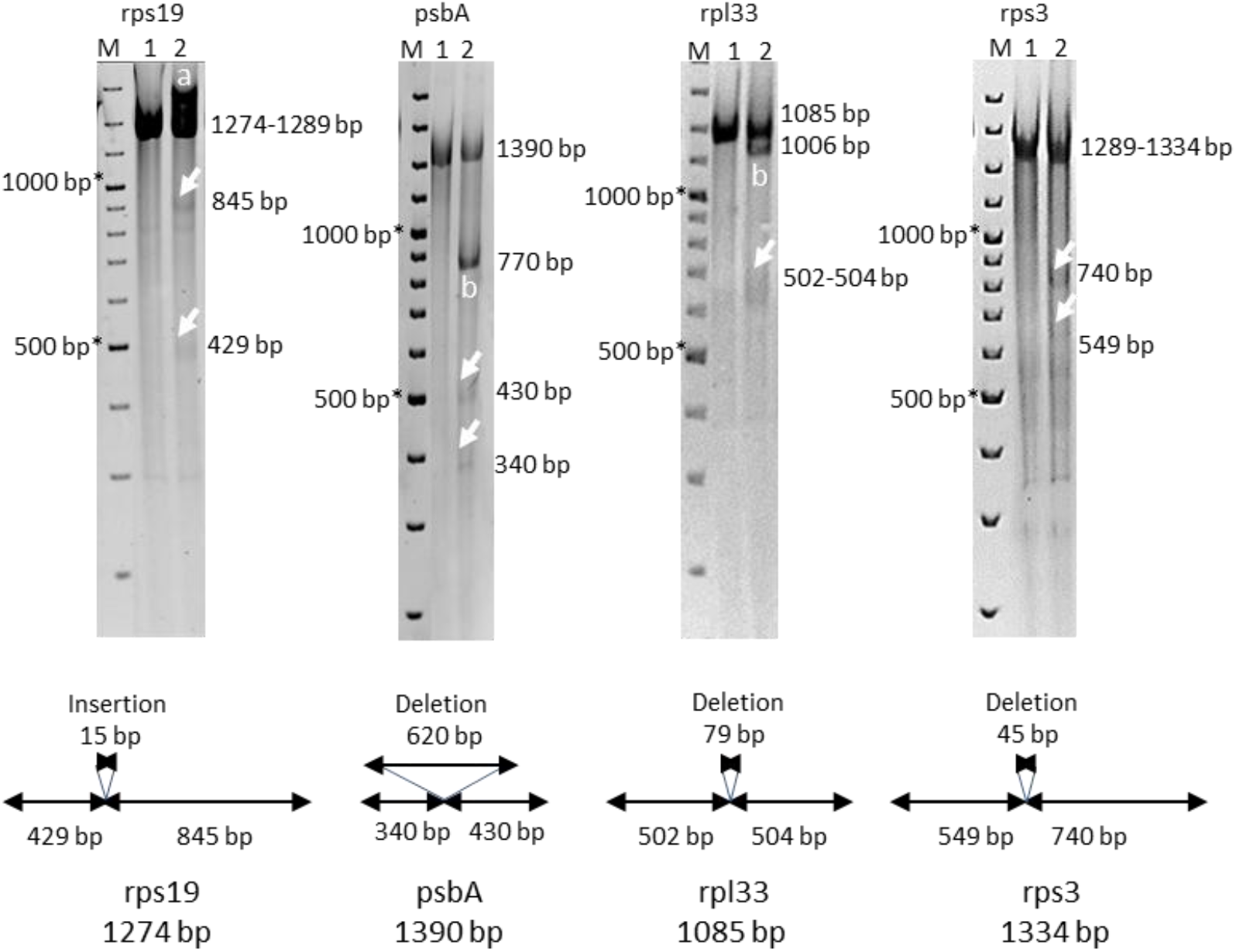
CJE digestions of amplicons carrying large indels: rps19, rpl33, psbA and rps3. M: molecular weight marker (100 bp ladder); 1: *cpm* seedling without the indels; 2: *cpm* seedling carrying large indel. Arrows point to the digestion bands; **a**: second band of amplification larger than the normal size of the amplicon; **b**: second band of amplification smaller than the normal size of the amplicon.*The samples run slower than the molecular weight marker probably due to the components of the CJE digestion mixture.

The larger band observed in the rps19 amplicon is hardly attributable to the 15 bp insertion segment since a 15 bp difference in size would not be resolved in a 3% polyacrylamide gel.

To investigate if larger bands similar to the one observed in the rps19 amplicon were also produced in the re-annealing step after the amplification of the other three amplicons and then digested with CJE, the amplicons were analyzed in 3% polyacrylamide gels, with or without CJE digestion (Fig. 2). The electrophoresis of all the amplicons without CJE digestion revealed a larger band than the expected size of the corresponding amplicon, which disappeared after CJE digestion. However, in the case of the rps19 amplicon, the digestion of this larger band was apparently incomplete, as also it seems to be in the original digestion in which the large indel was identified (see Figs. 1 and 2). Either the larger bands in the psbA and rpl33 amplicons were observed by mixing the *cpm* DNA with wild type DNA or by analyzing the *cpm* DNA samples alone, indicating the heteroplastomic state of these deletions. In contrast, for the indels in the rps19 and rps3 amplicons, the larger bands were only observed in the amplification of the *cpm* DNA samples mixed with wild type DNA, indicating the homoplastomic state of these indels.

**Figure 2.**
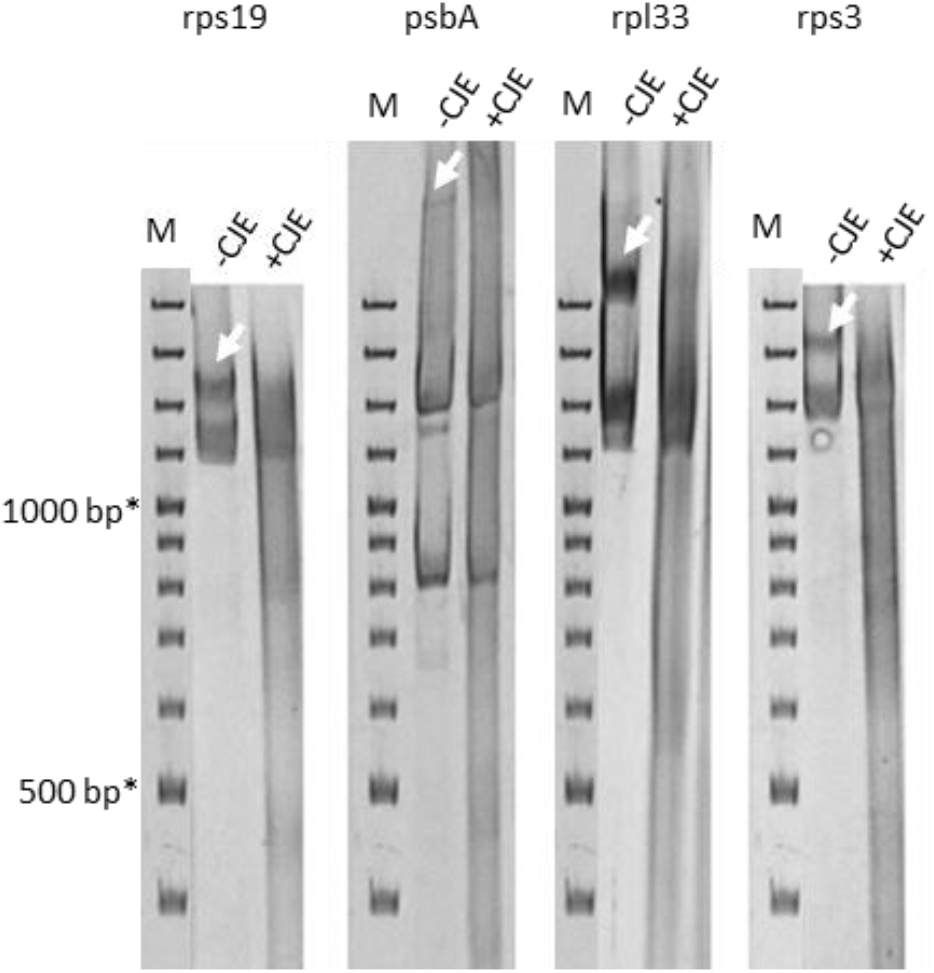
CJE digestions of the heteroduplexes coming from amplicons carrying large indels: rps19, psbA, rpl33 and rps3. Arrows indicate the presence of a band corresponding to the heteroduplexes before CJE digestion (-CJE), which disappears after CJE digestion (+CJE). M: molecular weight marker (100 bp ladder).*The samples run slower than the molecular weight marker probably due to the components of the CJE digestion mixture.

### Perfect or imperfect direct repeats were observed at both ends of the four large indels

Sequencing of the four amplicons carrying the large indels revealed the presence of short identical or similar sequences at both ends of the four large indels (see alignments in Fig. 3). These sequences were considered as perfect or imperfect direct repeats, respectively. Indels in the rps19 and psbA amplicons had imperfect direct repeats whereas indels in rpl33 and rps3 amplicons had perfect direct repeats. Bracketing the 15 bp tandem duplication in the *rps19 - tRNAHis* intergenic region, the direct repeats were not identical: one was 10 bp long, while the other one was nine bp long. The direct repeats also had a different size for the deletion in the *psbA* gene: one was 10 bp long and the other one was 17 bp long. In the case of the deletion in the intergenic region *rpl33 - rps18* and in the *rps3* gene, each pair of direct repeats were identical, consisting of seven and 24 bp, respectively. The sequences of the direct repeats are shown in Table 1.

**Figure 3.**
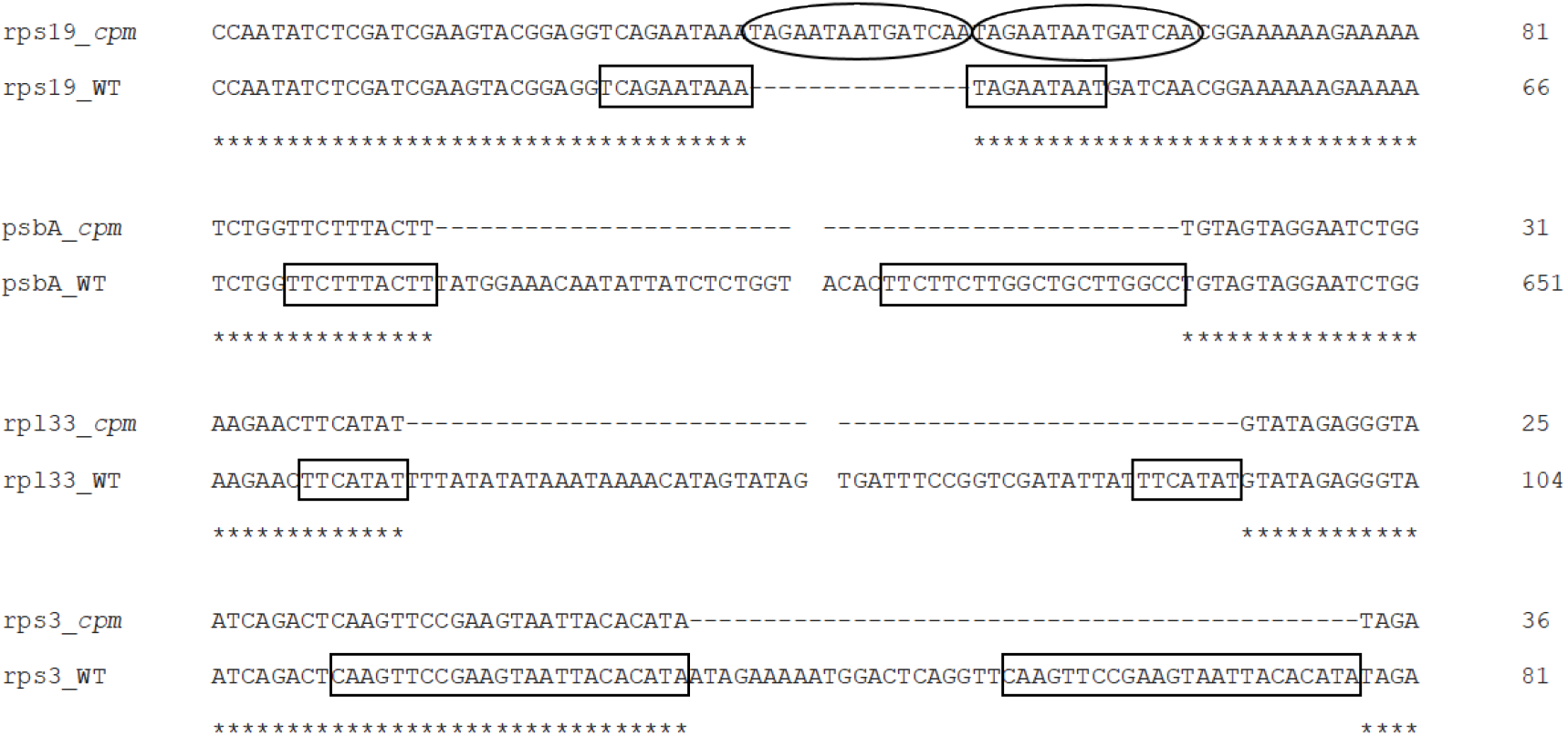
Alignments of amplicon segments flanking the large indels in *cpm* mutants and the sequences of wild type seedlings. Direct repeats are indicated in boxes and the duplicated sequence within the insertion of rps19 amplicon is shown by circles. Gaps in psbA and rpl33 amplicons indicate a discontinuous sequence.

**Table 1.**
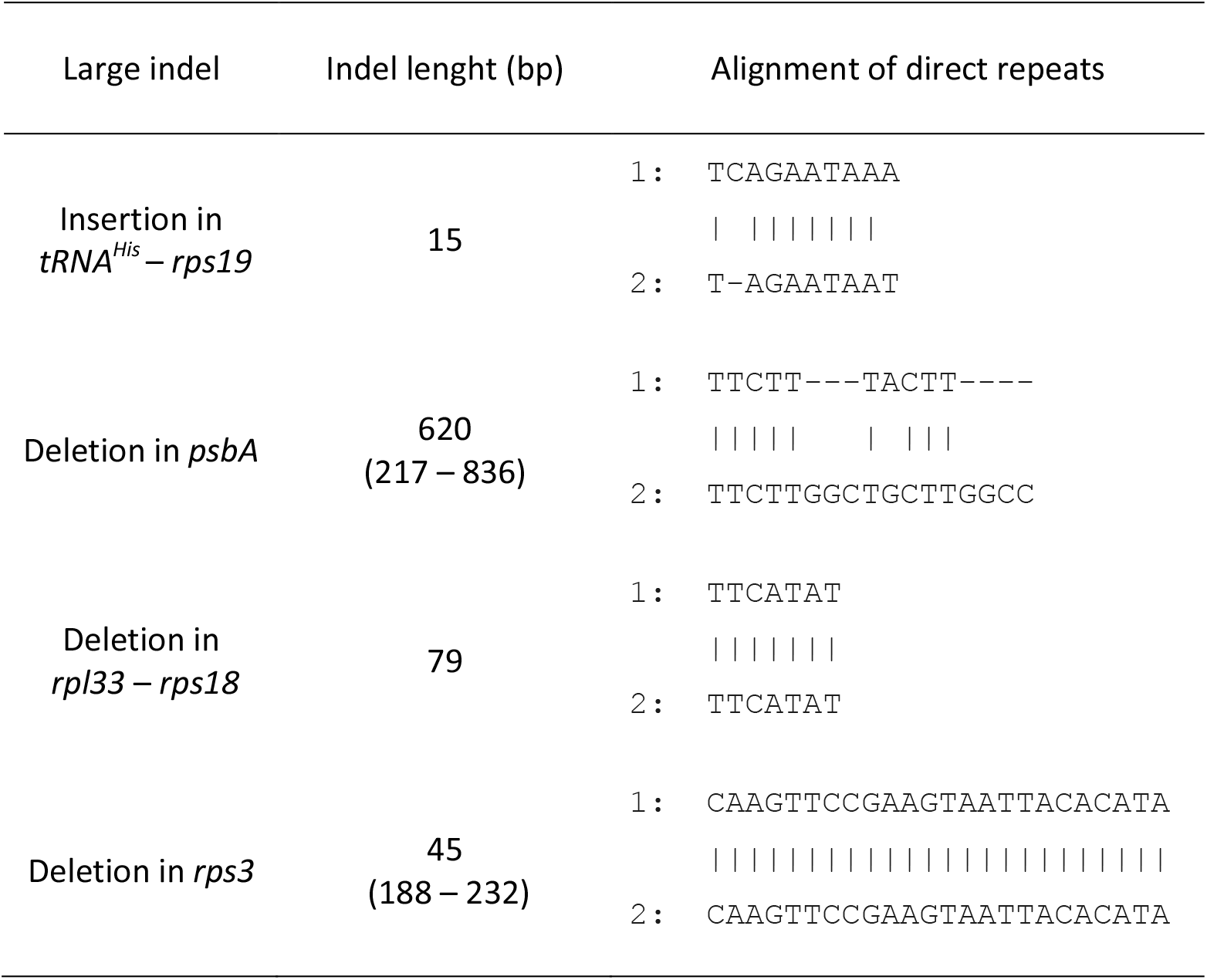
Direct repeats in large indels of rps19, psbA, rpl33 and rps3 amplicons. Numbers 1 and 2 correspond to the sequence of the repeat located to the left or to the right of the large indel, respectively. For the indels located in genes, the position of the beginning and end of the deleted bases are indicated in brackets according to the gene sequence. The insertion in *tRNA^His^ – rps19* intergenic region corresponds to duplication of a TAGAATAATGATCAA sequence.

### The protein sequence derived from the deleted version of the *rps3* gene of barley has the same size than the wild type RPS3 protein of maize

The 45 bp deletion found in the *rps3* gene does not produce a frameshift but eliminates a peptide of 15 amino acids in the RPS3 ribosomal protein of barley (Fig. 4). The *cpm* seedling carrying the deletion in homoplastomic state had normal green color but showed narrower and longer blades than the wild type control seedling.

**Figure 4.**
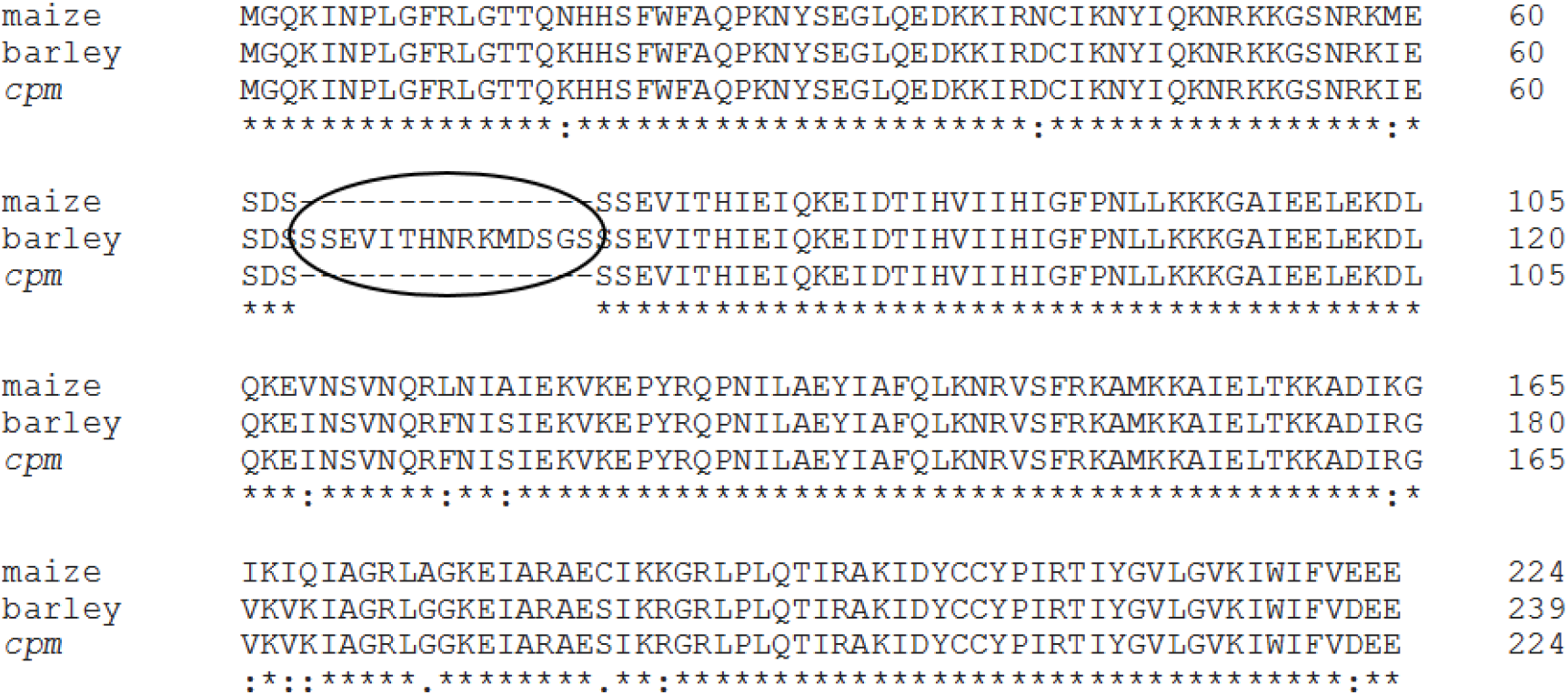
Alignment of the protein sequences of the *rps3* mutant isolated from *cpm* seedlings, the wild type sequence of barley and maize. The 15 amino acids lost by the deletion of 45 bp in the *cpm* seedling are indicated.

On the other hand, the alignment between the RPS3 polypeptide sequence of the *cpm* seedling carrying the deletion, the wild type version of barley and the orthologous RPS3 protein of maize revealed that the maize protein also lacks the segment eliminated in the *cpm* RPS3 protein (Fig. 4).

### The deletion in the *psbA* gene results in a non-functional photosystem II and causes seedling lethality

Originally, the deletion of 620 bp in the *psbA* gene was identified in heteroplastomy in one *cpm* seedling. Further screening of the progeny within the family from which the heteroplastomic seedling was identified allowed the isolation of one seedling carrying the same deletion in homoplastomy.

The 620 bp deletion in the *psbA* gene produces a frameshift and a premature stop codon, resulting in a truncated version of the D1 protein of 83 amino acids. The seedling carrying the homoplastomic deletion showed a *viridis* (homogeneous light green) phenotype and died two weeks after sowing.

To evaluate the effects of the truncated D1 protein, some thylakoid proteins were analyzed by Western blot in the *psbA* gene mutant and normal green *cpm* siblings. The *psbA* mutant showed aberrant expression of D1 and D2 proteins, both belonging to the photosystem II (PSII) core, whereas proteins of the photosystem I (PSI) core (PSI-A/B, D, E, L) and LHCB1 protein of the extrinsic antenna of PSII were expressed normally. In normal green *cpm* siblings, all these proteins were expressed (Fig. 5).

**Figure 5.**
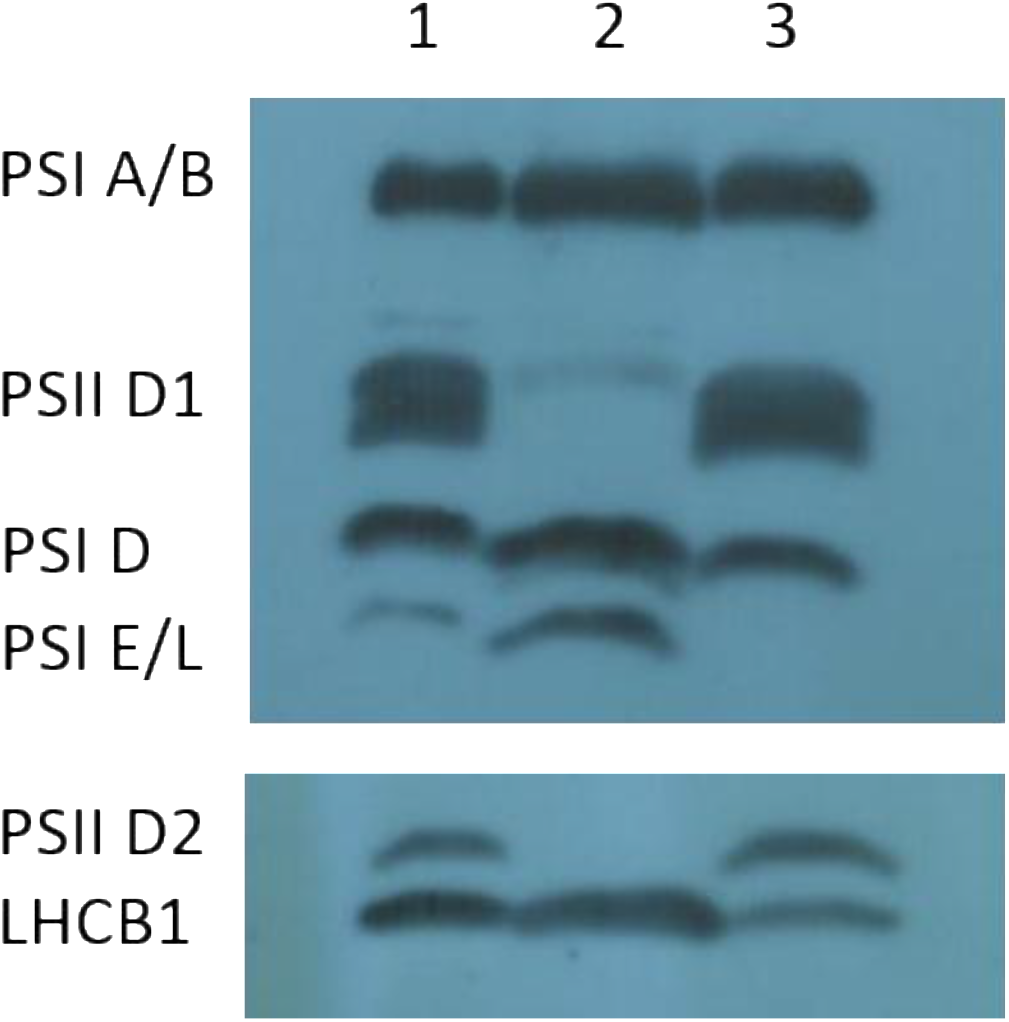
Immunoblot of photosystem proteins of the *viridis cpm* mutant homoplastomic for the 620 bp deletion in the *psbA* gene and in the *cpm* normal siblings. 1 and 3: normal green siblings. 2: *viridis* mutant.

## Discussion

The chloroplast genome is highly conserved in comparison with the plant nuclear genome, however, complex patterns of variation are observed when comparing different species (Clegg *et al*., 1994). Some differences correspond to length variations due to indels of several bases or larger (Ogihara *et al*., 1992). Extremely large indels have been observed at the junctions between the inverted repeats (IRs) and the large (LSC) or the small single copy regions (SSC) (Ogihara and Ohsawa, 2002), the intergenic region *16S rRNA-trnI* (Stoike and Sears, 1998), and the region between *rbcL* and *psaI* genes in grasses (Ogihara *et al*., 1988; Ogihara *et al*., 1991; Morton and Clegg, 1993; Ogihara and Ohsawa, 2002).

None of the four large indels investigated here were observed in the aforementioned regions, nor were they located in mononucleotide tandem repeats, in contrast to the many small indels (1 or 2 bp long) previously found by Landau *et al*. (2016). The finding of these large indels indicates that CJE can detect indels as large as 620 bp, which by far exceeds the size of the largest indel previously detected by TILLING in the nuclear genome (Comai *et al*., 2004). The digestion experiments demonstrated that CJE is able to digest the loop structure that would be generated during the heteroduplex formation by the mispairing of a wild type DNA strand with a strand carrying the indel, as postulated by Landau *et al*. (2016). An apparently incomplete digestion of the heteroduplex formed by the insertion in the rps19 amplicon was observed. This insertion was the smallest of the four large indels (15 bp) thus, a negative relationship between the digestion efficiency of CJE and the indel size could be the cause of the incomplete digestion. On the other hand, the larger size of the bands corresponding to heteroduplexes indicates that the loop structure affects the heteroduplex DNA mobility during electrophoresis. Other examples of decreased heteroduplexes mobility in polyacrylamide gels were shown by Fan *et al*. (2019).

It is worth noting that all the large indels studied here were flanked by short perfect or imperfect direct repeats (see Table 1). Recombination between short similar sequences (microhomologies), which can be considered a case of illegitimate recombination, has been suggested to be the cause of insertions and deletions in plastomes of grasses (Ogihara *et al*., 1998; Ogihara and Ohsawa, 2002). In nature, recombination between short similar repeats is highly controlled, which contributes to maintaining the integrity of the chloroplast genome and avoiding detrimental consequences of indels located in coding sequences. Consequently, this type of recombination event is rare and usually only observed on an evolutionary timescale (Maréchal and Brisson, 2010). The recombination process between repeats is less frequent when repeats are shorter, more distant, or when they carry more divergent sequences (Fischer *et al*., 1996). Our results support the hypothesis that illegitimate recombination between similar repeats flanking the deleted segments is the cause of the three large deletions detected in *cpm* seedlings. It has been reported that this type of recombination produces the loss of the DNA fragment between the repeats along with only one of the repeats (Ogihara *et al*., 1988, Ogihara *et al*., 1991; Ogihara and Osawa, 2002). The same situation was observed in the three deletions identified in *cpm* seedlings.

Another interpretation was offered for the large indels found in *Oenothera* plastome regions with many tandem direct repeats (Chang *et al*., 1996; Stoike and Sears, 1998). As seen in bacterial genomes (Bzymek and Lovett, 2001; Lovett, 2004), it was proposed that misalignment of direct repeats between the newly synthesized and template strands can generate indels through replication slippage. According to Lovett (2004), illegitimate recombination encompasses a number of distinct mechanisms, including slippage at short homologies as well as cut-and-join reactions at non-homologous sequences.

The complex patterns of plastome variability observed when comparing different species (Clegg *et al*., 1994) suggest occasional failures of the systems responsible for maintaining plastome genetic stability, in the otherwise highly conserved plastid genome (Zhang *et al*., 2016). Anti-recombination activity of MMR proteins can prevent illegitimate recombination between divergent short sequences by recognizing the mismatches present in the heteroduplexes of the intermediates of recombination (Lovett, 2004; Chakraborty and Alani, 2016). In fact, in the *msh1* gene (MMR family) mutant line of *A. thaliana*, in which anti-recombination activity of the MSH1 protein is deficient (Xu *et al*., 2011), some rearrangements (large indels) were identified in a plastome region carrying short similar repeats (10 - 15 bp).

We postulate that increased rates of illegitimate recombination could explain the occurrence of the large indels investigated here, taking place in *cpm* seedlings due to the absence of the anti-recombination activity of a MMR protein. Regarding this hypothesis, it is worth mentioning that the largest direct repeat of the rps3 amplicon is 24 bp, which is much smaller than 200 bp, a size that is usually mentioned for homologous recombination between repats in *E. coli* (Bi and Liu, 1994; Lovett 2004) and that is used for the incorporation of transgenes in the chloroplast genomes by homologous recombination.

With regard to the deleterious effects of the two large indels affecting coding sequences, two different situations were observed. In one case, the homoplastomic deletion in the *psbA* gene, which produces a frameshift and a premature stop codon, caused the loss of not only D1, but also D2 PSII core proteins, generating a nonfunctional PSII. This result is in agreement with observations in *C. reinhardtii*, where it is well established that these two proteins can accumulate only in the presence of each other. In the absence of D1 in the mutant FuD7, the D2 protein is synthesized but rapidly degraded (de Vitry *et al*., 1989). In contrast to the heterotrophic alga *C. reinhardtii*, for plants, the absence of D1 and D2 results in lethality (Järvi *et al*., 2015).

The other large indel affecting a coding region was the 45 bp deletion in the *rps3* gene, which eliminates a peptide of 15 amino acids. However, the mutant plant carrying the deletion in homoplastomy did not show a lethal effect: the seedling had a normal green color but the blades were narrower and longer than wild type barley. It has not yet been established if and how the altered morphology of the seedling is caused by the deletion affecting the *rps3* gene.

Interestingly, the 15 amino acids eliminated in the RPS3 protein of the barley mutant carrying the deletion in the *rps3* gene corresponds to a peptide that differentiates the wild type version of the RPS3 protein of maize and sorghum from the wild type version of barley (Saski *et al*., 2007). Analysis of the RPS3 protein sequence in grasses showed that these 15 amino acids are not always present (Saski *et al*., 2007), suggesting that this peptide is not necessary for the correct function of the protein.

## Conclusions

The presence of short perfect or imperfect direct repeats in the borders of the four large indels identified in the plastome of *cpm* seedlings is in agreement with the hypothesis that they originated as a consequence of increased rates of illegitimate recombination between these short repeats. Previously, illegitimate recombination was also postulated as the cause of the high frequencies of polymorphisms observed in the *rpl23* gene and pseudogene in *cpm* seedlings (Lencina *et al*., 2019). These observations together with the increased rates of substitutions and small indels in microsatellites previously observed in *cpm* seedlings (Landau *et al*., 2016) agree with the hypothesis of the malfunction of an MMR protein encoded by a nuclear gene in charge of maintaining plastome genetic stability in barley.

## Acknowledgements

We are very grateful to Prof. Barbara Sears for the invaluable suggestions on the original manuscript. We also would like to thank Mr. Abel Mario Moglie and Mr. José Cuello for skillful handling of the plant material.

